# Geomagnetic orientation in subterranean mole crickets during burrowing

**DOI:** 10.1101/2023.07.13.548821

**Authors:** Tsubasa Endo, Noriyasu Ando

**Affiliations:** Department of Life Engineering, Maebashi Institute of Technology, Gunma, Japan

**Keywords:** *Gryllotalpa orientalis*, magnetoreception, geomagnetic field, insects, orientation, burrowing

## Abstract

While many terrestrial animals use celestial and terrestrial cues for orientation, subterranean animals lack access to these cues. The geomagnetic field serves as a reliable alternative cue for underground navigation. Subterranean mammals use the geomagnetic field to burrow and navigate within their nests; however, whether subterranean insects possess magnetoreception remains unclear. Here, we investigated the influence of magnetic fields on burrowing directions in subterranean mole crickets (*Gryllotalpa orientalis*). Under an intact magnetic field, mole crickets predominantly burrow in the north-south direction. However, when the magnetic field was rotated by 90°, the preferred burrowing direction shifted to the east-west axis. Notably, when the horizontal component of the geomagnetic field was reduced to near 0 µT, no preferred direction was observed. These results suggest that mole crickets have a magnetic sense and they align the direction at the beginning of burrowing. The study enhances our comprehension of underground orientation mechanisms in subterranean insects and reveals convergent evolution between insects and mammals in utilizing the geomagnetic field as a navigational cue.

## 1. Introduction

Insects possess excellent navigation capabilities: they explore complex environments and localise to distant destinations using multiple sensory information. Vision (celestial and terrestrial cues) is essential for most terrestrial species to determine their direction [1]. They rely on the sun’s position and polarised light as well as the surrounding landscape to orient themselves [2]. In addition, they use non-visual cues such as odour, wind, and geomagnetic fields [3], to improve navigation accuracy and redundancy. Magnetoreception, the ability to detect the Earth’s geomagnetic field, serves as a global compass and is accessible regardless of time, weather, or light conditions. Magnetic sense has been reported in many invertebrate and vertebrates [4-6]. Among seasonally migratory insects, monarch butterflies employ a geomagnetic inclination compass as well as a sun compass [7, 8]. Migratory moths travelling at night also use a magnetic compass to calibrate their position relative to visual landmarks [9]. Magnetoreception is not limited to long-distance navigation; it is also utilised for specific tasks such as learning walk in ant navigation [10], constructing termite nests [11], and aligning the body presumably to determine direction [12]. However, the use of the magnetic compass is often secondary to vision or restricted to specific tasks in terrestrial insects that primarily rely on vision. On the other hand, the magnetic compass might be prioritised as an alternative to the sun compass in subterranean insects that burrow underground. In mammals, subterranean mole rats and their use of the geomagnetic compass have been extensively studied [13]. They begin constructing their nests at a specific azimuth and navigate inside the burrow using the magnetic compass [14, 15]. Furthermore, they refer to the axial information of the geomagnetic field when burrowing in straight paths [16]. While this phenomenon has been studied in subterranean mammals, it remains unclear whether subterranean insects use magnetoreception for underground orientation.

Here, we investigated oriental mole crickets (*Gryllotalpa orientalis*), which are non-migratory solitary insects that burrow nests and spend most of their life underground [17]. Therefore, the mole crickets offer a valuable demonstration of how subterranean mammals and insects have independently evolved similar strategies for navigating underground. We hypothesised that oriental mole crickets also have a magnetic sense and use it for burrowing behaviour. We investigated the influence of magnetic fields on their orientation by examining their burrowing trajectories when exposed to arbitrary magnetic fields.

## 2. Materials and methods

### (a) Experimental animals

Adult mole crickets (10 males and 19 females, Figure 1(a) and Supplementary Table S1) used in our experiment were kept solitary in a plastic cup with moistened sphagnum moss in our laboratory. They were maintained at a temperature of 25℃ under a 12:12 h light:dark cycle (Figure 1(a) and Supplementary Table S1).

**Figure 1.**
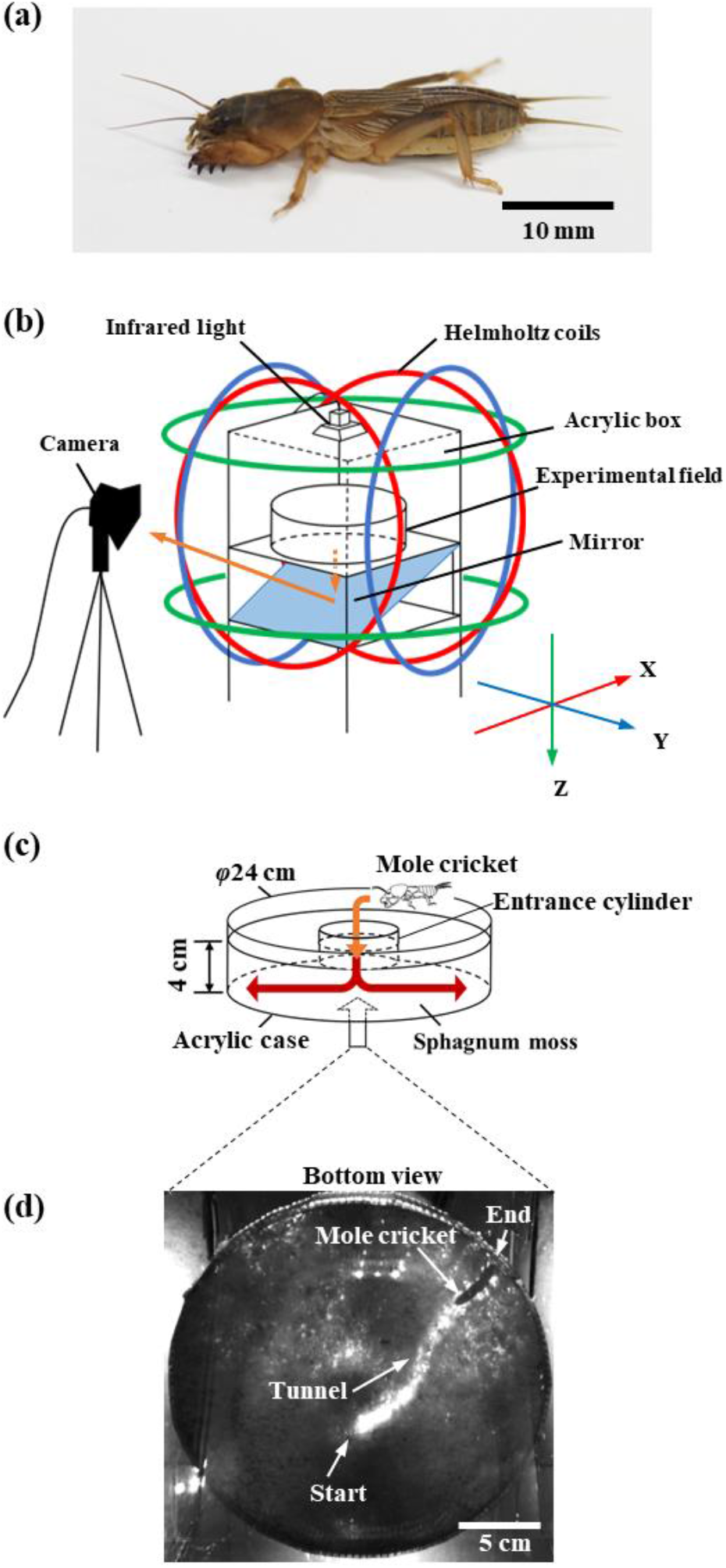
Experimental setup. (a) Mole cricket. (b) Experimental system. The magnetic field was generated using Helmholtz coil in three directions (X, Y, and Z). A mirror was placed under the experimental field to capture the behaviour of the mole crickets with an infrared camera. (c) Experimental field. The mole crickets burrowed through the sphagnum moss vertically from inside the entrance cylinder (orange vertical arrow) and toward the sides of the field when they reached the bottom (red horizontal arrows). (d) Bottom view of the experimental field when a mole cricket reached the end of the field, captured by the infrared camera.

### (b) Experimental system and magnetic conditions

An experimental system was designed to analyse the burrowing behaviour of mole crickets in a horizontal plane when exposed to different magnetic fields (Figure 1(b)). The magnetic field was generated using a Helmholtz coil consisting of a pair of circular coils (diameter: 60–68 cm) which were installed along three axes (X-, Y-, and Z-axes). Direct current was applied to each coil to generate a magnetic field of arbitrary strength and direction inside the coil system. An acrylic box (30 × 30 × 30 cm) was placed inside the coil system (50 cm from the floor), and as the experimental field, a cylindrical acrylic case (diameter: 24 cm, height: 6 cm; Figure 1(c)) was placed at the centre of the acrylic box. The field was filled with moistened sphagnum moss (depth: 4 cm). Infrared light (wavelength: 850 nm) was placed above the box to serve as a background light to capture the burrowing behaviour from the bottom.

Three horizontal magnetic conditions were set: the magnetic field in the laboratory (LMF, horizontal intensity of 41.7 μT), a 90°-rotated magnetic field (RMF, 41.8 μT), and a zero-magnetic field (ZMF, with a horizontal intensity of nearly zero: 0.11 μT). The coils of the X- and Y-axes were activated to generate a rotated magnetic field in the RMF and to cancel the LMF in the ZMF, while they were inactivated in the LMF. The inclination angle of the experimental condition was shallow because of the small Z-component in the laboratory (inclination angle/Z-component, 11.7°/8.6 μT in the laboratory vs 50.4°/36.0 μT outside). A supplementary experiment was conducted to investigate the influence of the inclination angle by activating the Z-axis coil (Supplementary Figure S4). Detailed specifications of the magnetic conditions are provided in Supplementary Table S2 and Figure S1.

### (c) Capturing of burrowing behaviour

When released into an entrance cylinder (φ = 60 mm) partially stuffed at the experimental field centre (depth: 20 mm), the mole crickets initiated their burrowing behaviour by starting to burrow vertically. Subsequently, they transitioned to burrowing horizontally at the bottom of the field (see Figure 1(c)). The release direction (either north, south, east, or west of the LMF) was randomised for each trial. Mole crickets burrowing horizontally were reflected on a mirror below the experimental field (Figure 1(d)), and their burrowing behaviour was captured every second using an infrared camera (DN3V-200BU; Shodensha, Osaka, Japan). The burrowing trajectories of 24 individuals were studied and captured under each magnetic field condition, with three trials conducted for each individual (total of 72 trials). All experiments were conducted in a dark room during the dark period, and the sides of the experimental field and acrylic box were covered with black paper to eliminate the influence of visual cues on their behaviour. Sphagnum moss was stirred in each trial and moss was replaced for each individual to avoid possible effects of olfactory or mechanical biases in the experimental field.

### (d) Data analysis

The horizontal burrowing trajectories were captured by tracking the mole crickets’ heads. The position at which they first reached the bottom after entering the field was defined as the origin. The majority of trajectories were straight from the origin to the wall; however, as some were curved, the body axis angle was obtained at three different positions to serve as orientation indices: 1.5 cm from the origin (initial position); 6 cm from the origin (intermediate position); and where the insect reached the wall (endpoint). Because the preliminary observations of body angles revealed a bimodal distribution, they were treated as axial data, and a mean vector (*θ*, mean axial direction; *r*, mean vector length) was calculated for each magnetic condition. The Rayleigh test was used to determine significant deviations from a random distribution, and Watson’s two-sample test was used to test for homogeneity in two angular distributions (significance level, *P* < 0.05). For details, see Supplementary Figures S2 and S3.

## 3. Results

The duration of a single trajectory (from beginning to end of burrowing) are presented as median (interquartile range) and were as follows: LMF: 102.0 (59.8–165.8) s, RMF: 89.5 (54.8–140.5) s, and ZMF: 85.5 (52.5–120.0) s; and there was no significant difference in the burrowing duration between magnetic conditions (*P* >0.05, Steel-Dwass test). The overall burrowing trajectories in the LMF showed a bimodal distribution in the north-south direction, whereas in the RMF, they were distributed in the east-west direction. However, in the ZMF, no remarkable tendency in any specific direction was observed (Figure 2(a)). Using the Rayleigh test, significant deviations from random orientation was detected at 1.5 cm (*θ* = 10° / 190°, *P* = 0.002) and 6 cm (*θ* = 15° / 195°, *P* = 0.018) in the LMF (Figure 2(b)), indicating that the orientation was biased toward the north-south axis. Significant deviations from random orientation were also found at 6 cm (*θ* = 66° / 246°, *P* = 0.02) and endpoints (*θ* = 73° / 253°, *P* = 0.049) in the RMF, indicating that the orientation was biased approximately toward the east-west axis (Figure 2(c)). Comparison of the angular distributions between the LMF and RMF revealed a significant difference at 6 cm (*P* = 0.004, Watson’s two-sample test). In contrast, no significant orientation bias was observed in the ZMF (Figure 2(d)).

**Figure 2.**
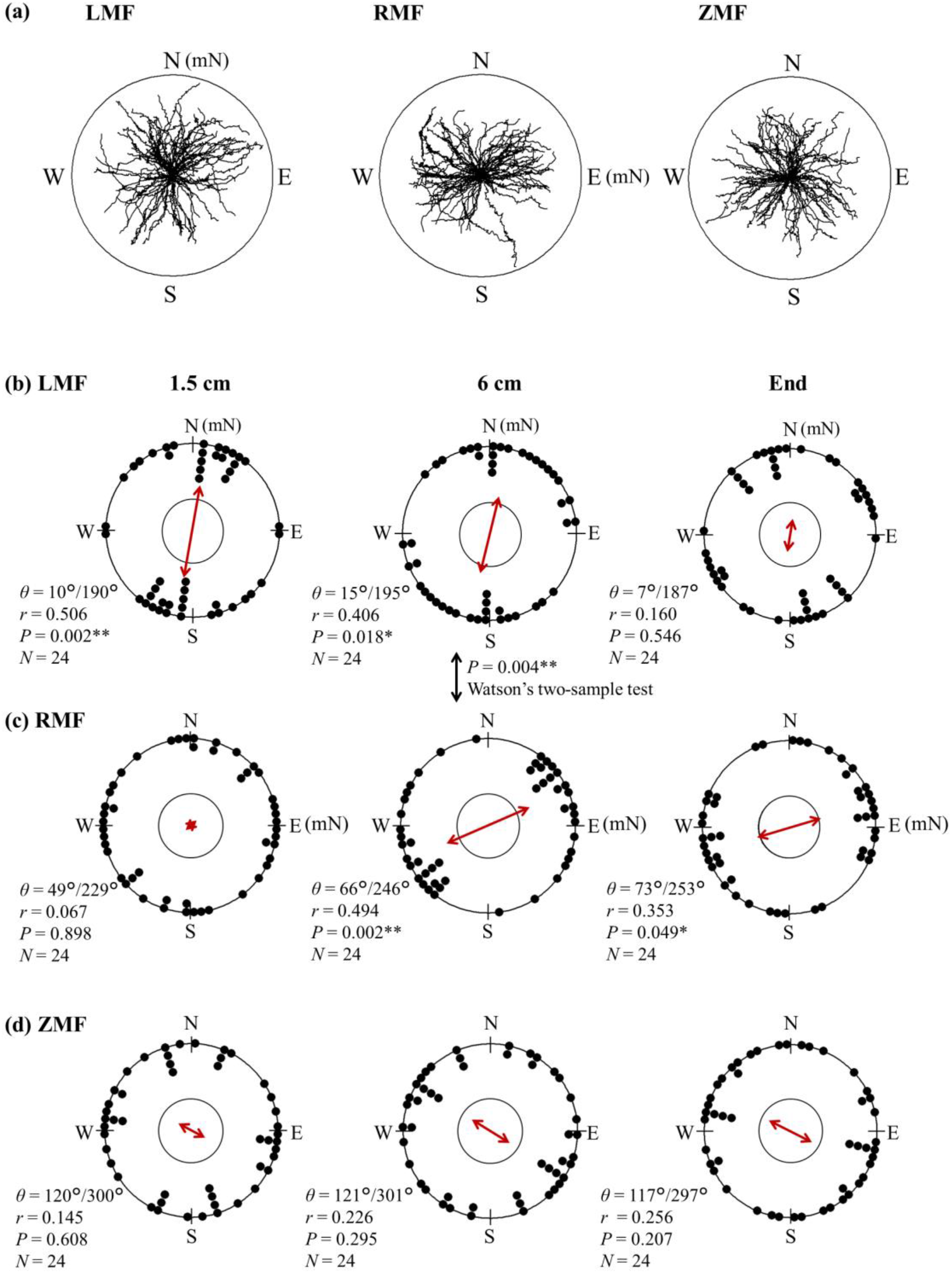
Orientation of burrowing behaviour. (a) Trajectories of the 72 trials of 24 individuals under three magnetic conditions (magnetic field in the laboratory (LMF), 90°-rotated magnetic field (RMF), and zero-magnetic field (ZMF)). (b–d) Body angle orientation at three points (1.5 cm and 6 cm from the origin, and at the endpoint) under the three magnetic conditions. The mean axial vector is indicated by a red two-way arrow with a radius ranging from 0 to 1. A single black point on the circumference represents the mean body angle of one individual (three trials), plotted as axial data on a circumference of radius 1. The inner circle represents the *P* = 0.05 significance level calculated using the Rayleigh test. *θ* is the angle of the mean axial vector when the magnetic north in LMF is 0°, *r* is the length of the mean axial vector, *P* is the *P*-value calculated with the Rayleigh test, and *N* is the number of individuals subjected to the trial. N, E, S, and W around the circumference represent north, south, east, and west in the laboratory (LMF), respectively. mN represents magnetic north. Watson’s two-sample test was used to compare the distributions of body orientations between two magnetic conditions. Asterisks indicate statistical significance (*P* < 0.05*, < 0.01**).

## 4. Discussion

Our results showed a tendency in mole crickets to align their body angles with the magnetic north-south axis. Possible experimental biases that influence burrowing direction are vision, entry direction, substrate (sphagnum moss) and olfaction. As our experiments were performed in the dark, the visual bias was eliminated. By randomising the direction of release in each trial, stirring and replacing the moss between trials and individuals, respectively, and keeping the position of the experimental equipment unchanged under all magnetic conditions, further possible biases were minimised. Therefore, our results suggested that the observed orientation tendency depended on the magnetic field and that the mole cricket has magnetic sense. The observed bimodal body angle distribution suggested that *G. orientalis* exhibits magnetic alignment, similar to various animal species [12].

The difference in mean vector length between the LMF and RMF at 1.5 cm was noticeable (Figure 2(b, c)). Because the studied insects in the RMF experienced a change in the horizontal magnetic field when placed in the RMF, we speculated that they may have required some time to adapt and align to the new magnetic field. A recent study reported that the night-migratory bogong moth uses a magnetic field for migration, however, there is a delay in flight direction changes after the magnetic field direction is altered. This suggest that the reference to the magnetic field is periodic rather than continuous [9].

The possible influence of the shallow inclination angle in the LMF (an inclination angle of 11.7°, corresponding to the inclination near the equator) should also be noted. We conducted an additional experiment in which we reproduced the measured geomagnetic conditions in our institute at ground level (X, Y, Z = 29.3, 0.03, 36.0 μT; 36°21’ N, 139°04’ E) and confirmed that the mole crickets also showed significant distribution biases of body angle toward the north-south magnetic axis in all three measurement points (Supplementary Figure S4). Although we did not completely cancel the Z component, the bimodal distribution of the body angle at the two inclination angles suggests that mole clickets have a declination compass rather than an inclination compass.

*G. orientalis* constructs vertical and horizontal burrows for foraging. They use vertical burrows to hide from predators, rest, moult, and overwinter, and horizontal burrows to escape and mate [17]. Horizontal burrowing is shallow (< 5 cm deep) and spreads with branches, with its pattern influenced by food distribution [18]. A mole cricket’s underground orientation during burrow construction is important for efficient foraging as well as for successful escape, as escape routes in burrows are limited. In particular, burrowing in a straight line is optimal for expanding the burrow area and moving quickly inside the tunnel at the shortest distance between two points; however, it requires external sensory references [19]. In mammals, subterranean rodents can burrow long, straight tunnels aligned at a certain angle relative to the direction of the geomagnetic field [20]. The horizontal burrowing of *G. orientalis* also follows relatively straight paths [17, 18]; therefore, our results suggest that these insects and subterranean mammals share a strategy of burrowing in a straight line with reference to the geomagnetic field. On the other hand, it should be noted that our analysis was limited to a specific phase of burrow construction. We analysed the behaviour from the beginning of horizontal burrowing, and all trajectories were single tracks with no branches (Figure 2(a)), suggesting that the behaviour represented an early phase of horizontal burrowing during which unknown environments would be explored. A recent study reported that hunting dogs mostly start with a short run along the north-south geomagnetic axis in the inbound track using novel routes (scouting) [21]. Although the burrowing in this study is not a homing behaviour, a straight movement with reference to the geomagnetic field is an effective strategy for exploring unknown environments.

Two models have been proposed to describe the mechanism of magnetoreception: photochemical reaction of cryptochromes (photoreceptive proteins), and the mechanical force of magnetic substances (magnetite) present *in vivo* [22]. Considering that mole crickets are nocturnal and burrow underground, and that our experiments were conducted in the dark, it is likely that they rely on a magnetite-based mechanism for underground orientation, similar to subterranean mole rats [23]. However, the possibility of using cryptochromes cannot be ruled out because they also play a role in the circadian clock, and animals need to calibrate the clock using light. Further analysis of compass types (declination or inclination) and magnetoreceptive organs should be carried out to describe the mechanism of magnetoreception in mole crickets, providing a basis for comparison with subterranean mammals as an example of convergent evolution optimised for underground orientation.

## Ethics statement

The animal experiments were performed in accordance with the guidelines of the Maebashi Institute of Technology.

## Data accessibility

The datasets supporting this article have been uploaded as part of the supplementary material.

## Authors’ contributions

T. E.: conceptualisation, methodology, investigation, formal analysis, writing, and editing; N.A.: conceptualisation, methodology, writing, and editing.

## References

[1] Gould, J.L. 1998 Sensory bases of navigation. Curr Biol 8, R731–R738. (doi:https://doi.org/10.1016/S0960-9822(98)70461-0).

[2] Wolf, H. 2011 Odometry and insect navigation. J Exp Biol 214, 1629–1641. (doi:10.1242/jeb.038570).

[3] Heinze, S. 2017 Unraveling the neural basis of insect navigation. Curr Opin Insect Sci 24, 58–67. (doi:10.1016/j.cois.2017.09.001).

[4] Vácha, M. 2020 Invertebrate magnetoreception – In between orientation and general sensitivity. In Reference Module in Neuroscience and Biobehavioral Psychology (ed. B. Fritzsch), pp. 445–458. Oxford, Elsevier.

[5] Wiltschko, W. & Wiltschko, R. 2005 Magnetic orientation and magnetoreception in birds and other animals. J Comp Physiology A 191, 675–693. (doi:10.1007/s00359-005-0627-7).

[6] Wiltschko, R. & Wiltschko, W. 2022 The discovery of the use of magnetic navigational information. J Comp Physiol A 208, 9–18. (doi:10.1007/s00359-021-01507-0).

[7] Guerra, P.A., Gegear, R.J. & Reppert, S.M. 2014 A magnetic compass aids monarch butterfly migration. Nat Commun 5, 4164. (doi:10.1038/ncomms5164).

[8] Reppert, S.M., Guerra, P.A. & Merlin, C. 2016 Neurobiology of monarch butterfly migration. Annu Rev Entomol 61, 25–42. (doi:10.1146/annurev-ento-010814-020855).

[9] Dreyer, D., Frost, B., Mouritsen, H., Gunther, A., Green, K., Whitehouse, M., Johnsen, S., Heinze, S. & Warrant, E. 2018 The Earth’s magnetic field and visual landmarks steer migratory flight behavior in the nocturnal Australian bogong moth. Curr Biol 28, 2160–2166 e2165. (doi:10.1016/j.cub.2018.05.030).

[10] Fleischmann, P.N., Grob, R. & Rössler, W. 2020 Magnetoreception in Hymenoptera: importance for navigation. Animal Cognition 23, 1051–1061. (doi:10.1007/s10071-020-01431-x).

[11] Jacklyn, P. & Munro, U. 2002 Evidence for the use of magnetic cues in mound construction by the termite Amitermes meridionalis (Isoptera : Termitinae). Australian Journal of Zoology 50. (doi:10.1071/ZO01061).

[12] Begall, S., Malkemper, E.P., Červený, J., Němec, P. & Burda, H. 2013 Magnetic alignment in mammals and other animals. Mammalian Biology 78, 10–20. (doi:10.1016/j.mambio.2012.05.005).

[13] Begall, S., Burda, H. & Malkemper, P. 2014 Magnetoreception in mammals. Adv Stud Behav 46, 45–88. (doi:10.1016/B978-0-12-800286-5.00002-X).

[14] Burda, H., Marhold, S., Westenberger, T., Wiltschko, R. & Wiltschko, W. 1990 Magnetic compass orientation in the subterranean rodent Cryptomys hottentotus (Bathyergidae). Experientia 46, 528–530. (doi:10.1007/bf01954256).

[15] Kimchi, T., Etienne, A.S. & Terkel, J. 2004 A subterranean mammal uses the magnetic compass for path integration. Proc Natl Acad Sci USA 101, 1105. (doi:10.1073/pnas.0307560100).

[16] Malewski, S., Begall, S., Schleich, C.E., Antenucci, C.D. & Burda, H. 2018 Do subterranean mammals use the Earth’s magnetic field as a heading indicator to dig straight tunnels? PeerJ 6, e5819. (doi:10.7717/peerj.5819).

[17] Endo, C. 2007 The underground life of the oriental mole cricket: an analysis of burrow morphology. J Zool 273, 414–420. (doi:https://doi.org/10.1111/j.1469-7998.2007.00345.x).

[18] Endo, C. 2008 An analysis of the horizontal burrow morphology of the oriental mole cricket (Gryllotalpa orientalis) and the distribution pattern of surface vegetation. Can J Zoolog 86, 1299–1306. (doi:10.1139/z08-116).

[19] Cheung, A., Zhang, S., Stricker, C. & Srinivasan, M.V. 2007 Animal navigation: the difficulty of moving in a straight line. Biol Cybern 97, 47–61. (doi:10.1007/s00422-007-0158-0).

[20] Eloff, G. 1951 Orientation in the mole-rat Cryptomys. British Journal of Psychology. General Section 42, 134–145. (doi:10.1111/j.2044-8295.1951.tb00285.x).

[21] Benediktova, K., Adamkova, J., Svoboda, J., Painter, M.S., Bartos, L., Novakova, P., Vynikalova, L., Hart, V., Phillips, J. & Burda, H. 2020 Magnetic alignment enhances homing efficiency of hunting dogs. Elife 9. (doi:10.7554/eLife.55080).

[22] Byrne, J.H. & Vacha, M. 2019 Magnetoreception of invertebrates. In The Oxford Handbook of Invertebrate Neurobiology (pp. 366–388.

[23] Caspar, K.R., Moldenhauer, K., Moritz, R.E., Nemec, P., Malkemper, E.P. & Begall, S. 2020 Eyes are essential for magnetoreception in a mammal. J R Soc Interface 17, 20200513. (doi:10.1098/rsif.2020.0513).

