## Supplementary materials for "Geomagnetic orientation in subterranean mole crickets during burrowing"

#### 1. Insects

Oriental mole crickets (*Gryllotalpa orientalis*) were purchased from private collectors, and 29 adult mole crickets were used in the experiment (Table S1). To rear the mole crickets, one plastic cup (129 mm diameter × 97 mm height) filled with sphagnum moss was used per individual. All rearing cups were maintained in an incubator (CN-40A; Mitsubishi Electric Engineering Corporation, Tokyo, Japan). The incubator was kept at 25 °C, a 12-hour light period from 0:00 to 12:00, and a 12-hour dark period from 12:00 to 24:00. All experiments were conducted during the dark period because mole crickets are nocturnal insects.

**Table S1** Details of the mole crickets used in the experiment.

| ID | Length [cm] | Weight [g] | Sex | LMF | RMF | ZMF |
|---|---|---|---|---|---|---|
| 1 | 2.7 | 0.86 | female | ● | ● | |
| 2 | 2.6 | 0.68 | male | ● | ● | ● |
| 3 | 2.5 | 0.91 | female | ● | ● | ● |
| 4 | 2.9 | 0.91 | female | ● | ● | ● |
| 5 | 2.8 | 0.99 | female | ● | ● | ● |
| 6 | 2.7 | 0.83 | female | ● | ● | |
| 7 | 2.8 | 0.90 | female | ● | ● | ● |
| 8 | 2.9 | 1.05 | female | ● | ● | ● |
| 9 | 2.8 | 1.08 | female | ● | ● | ● |
| 10 | 2.4 | 0.69 | male | ● | ● | |
| 11 | 2.9 | 1.01 | male | ● | ● | |
| 12 | 2.5 | 0.66 | male | ● | ● | |
| 13 | 2.6 | 0.77 | female | ● | ● | ● |
| 14 | 2.7 | 0.63 | female | ● | ● | ● |
| 15 | 3.1 | 1.17 | female | ● | ● | ● |
| 16 | 2.5 | 0.71 | female | ● | ● | ● |
| 17 | 3.1 | 0.94 | female | ● | ● | ● |
| 18 | 2.8 | 0.78 | female | ● | ● | ● |
| 19 | 2.8 | 0.95 | male | ● | ● | ● |
| 20 | 3.0 | 0.94 | male | ● | ● | ● |
| 21 | 3.0 | 1.10 | female | ● | ● | ● |
| 22 | 2.9 | 1.10 | male | ● | ● | ● |
| 23 | 3.0 | 0.87 | male | ● | ● | ● |
| 24 | 2.4 | 0.66 | female | ● | ● | ● |
| 25 | 2.4 | 0.55 | female | | | ● |
| 26 | 2.2 | 0.37 | male | | | ● |
| 27 | 2.8 | 0.89 | female | | | ● |
| 28 | 2.9 | 0.93 | male | | | ● |
| 29 | 2.6 | 0.74 | female | | | ● |

Black circles (●) indicate which magnetic conditions were used for each individual. LMF: magnetic field in the laboratory, RMF: 90°-rotated magnetic field, ZMF: nearly zero magnetic field.

### 2. Magnetic condition

#### 2.1 Magnetic field parameters

The parameters of the three magnetic conditions (magnetic field in the laboratory (LMF), 90°-rotated magnetic field (RMF), and nearly zero magnetic field (ZMF)) used in the experiment are listed in Table S2. The parameters were based on the measured central magnetic field of an acrylic box (0.3 m × 0.3 m) placed inside the coil. In the table, X, Y, and Z represent the component intensities of the magnetic field measured using a magnetic sensor (BM1422AGMV; ROHM, Kyoto, Japan). The X-axis direction corresponds to the magnetic field north of the LMF. The declination angle represents the angular deviation of the magnetic field from the geographic north, and the geomagnetic field at Maebashi Institute of Technology (Maebashi City, Gunma Prefecture, 36°21'N, 139°04'E), where the experiment was conducted, shifted -7.67° from the geographic north, which was almost the same as that in the laboratory (-7.76°, LMF). Therefore, in this experiment, magnetic north (N) in LMF was defined as the orientation of 7.76° from the geographic north.

The RMF was a 90° clockwise rotation of the horizontal components of the LMF in which the X and Y components of the magnetic field were manipulated. In addition, the ZMF cancelled the X and Y components of the magnetic field. However, the Z-direction (vertically downward) of the magnetic field remained. All the experiments were conducted in the laboratory, with a difference in the angle of inclination (I) between the magnetic field measured in the LMF and the geomagnetic field measured on the ground located near the laboratory (GMF, Table S2). All mole crickets in this experiment were reared and exposed to a geomagnetic field in the laboratory; therefore, the LMF represents the magnetic field in the laboratory without any manipulation. To investigate the influence of a shallow inclination angle in the laboratory, we also measured the burrowing trajectories of mole crickets under magnetic conditions equal to the geomagnetic field (Figure S3).

**Table S2** Magnetic field parameters for the burrowing experiments.

| | X(μT) | Y(μT) | Z(μT) | F(μT) | D(°) | I(°) | H(μT) |
|---|---|---|---|---|---|---|---|
| LMF | 41.67 | 0.062 | 8.642 | 42.56 | -7.756 | 11.72 | 41.67 |
| RMF | -0.173 | 41.84 | 8.409 | 42.68 | 82.57 | 11.36 | 41.84 |
| NMF | -0.117 | -0.011 | 8.595 | | | | 0.117 |
| Geomagnetic field (ground) | 29.29 | 0.029 | 35.97 | 46.59 | -7.612 | 50.44 | 29.68 |

The magnetic parameters refer to the value of the central magnetic field of the acrylic box (0.3 × 0.3× 0.3 m) placed inside the coil. X, Y, Z: Magnetic components measured using magnetic sensors. X represents the magnetic north in the LMF, while Z represents the vertical direction and is positive downward. F: total intensity; D: Declination; I, Inclination; H: Horizontal intensity.

### 2.2 Distribution of magnetic parameters

The magnetic parameters (total intensity, declination, inclination, and horizontal intensity) within the acrylic box placed inside the coil are shown in a heat map (Figure S1). The squares with 30 cm long sides in the figure represent the bottom of the acrylic box, and the inner circle represents the experimental field (24 cm in diameter). The distribution of the parameters in the acrylic box is represented by different colours, and the barometer on the right represents the colour-coded parameter values. One side of the acrylic box (X) was placed parallel to the magnetic north (mN) in the laboratory. The magnetic field was generated by adjusting the acrylic box such that its centre became the target value. Although there were some variations in the intensity and directional distribution of the generated magnetic field (especially at the edges away from the centre of the acrylic box), the variation was relatively small compared to the target intensity and direction, suggesting that the effects of the variation in the magnetic field generated within the experimental field on the behaviour of the mole crickets can be expected to be small.

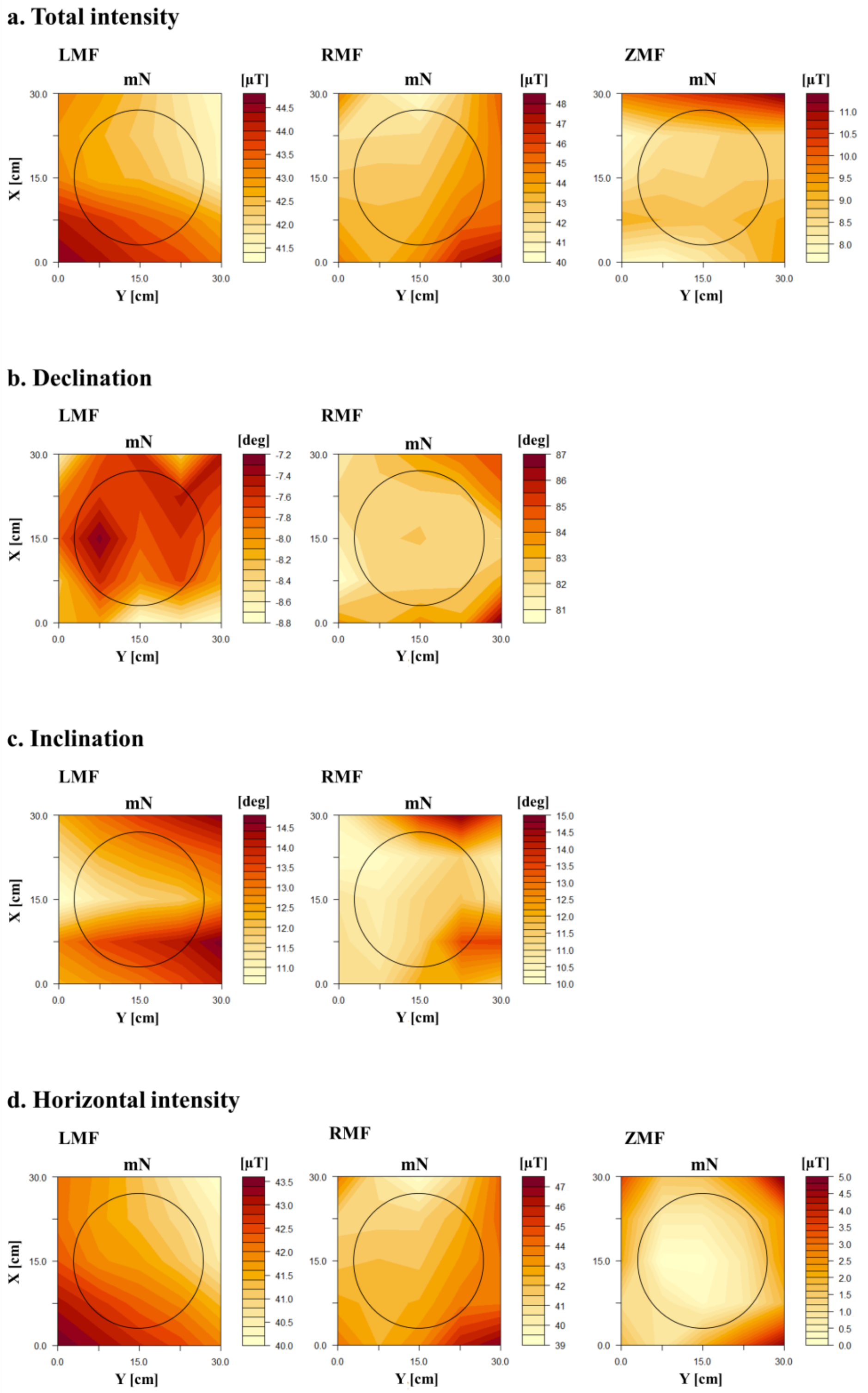


v

**Figure S1** Heat map showing the distribution of the magnetic parameters (total intensity, declination, inclination, and horizontal intensity) within the acrylic box placed inside the coil map. The inner circle represents the experimental field (24 cm in diameter). mN, magnetic north.

### 3. Data analysis

#### 3.1 Burrowing trajectory

The burrowing trajectories of the mole crickets were calculated by analysing the images at the bottom of the experimental field. The starting point of the measurement was when the mid-legs of the mole crickets were visible, and the endpoint was when the mole crickets reached the edge of the experimental field. Using a video analysis software (DIPP-Motion V Ver1.2.5, Ditect, Tokyo, Japan), the coordinates of the tips of the mole crickets' heads were measured at 1-second intervals (Figure S2), and the burrowing trajectory was determined. All results are shown with the starting point aligned to the origin.

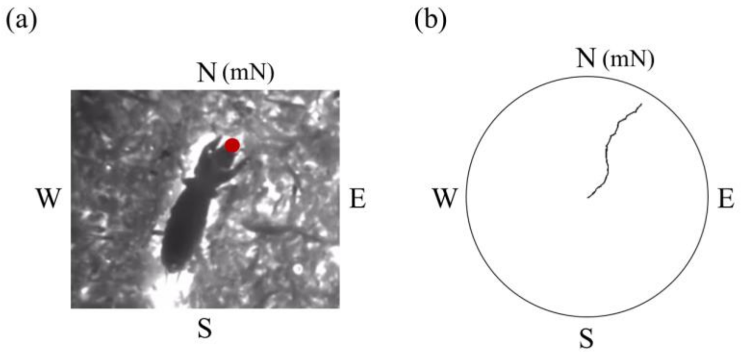


**Figure S2** Burrowing trajectories. (a) Photograph of a mole cricket. The mole crickets digging at the bottom of the field were captured using a mirror. (b) Burrowing trajectory of one individual. In both (a) and (b), N, E, S, and W around the circumference represent magnetic north, south, east, and west in the laboratory (LMF), respectively. mN represents magnetic north.

### 3.2 Analysis of body orientation

To determine the burrowing direction for all mole crickets (24 individuals with three trials each, a total of 72 trials in each magnetic condition), the mean axial vector was calculated under the three magnetic conditions. The analysis was performed at 1.5 cm, 6 cm, and endpoint from the starting point of the burrowing trajectory. As an example, the procedure for calculating the mean axial vector at 6 cm in the geomagnetic field was as follows:

(a) Extraction of body angles (Figure S3(a)): The body angles of the mole crickets at 6 cm were calculated from their burrowing trajectories. The body angle was defined as the angle (0° = north in LMF (N)) of the line connecting the tip of the head (red) and the middle of the mid-leg (green) from the image of the mole crickets taken at the bottom of the experimental field. Because the burrowing trajectory obtained in this experiment was bimodal along the magnetic axis, the body angle was treated as axial data for the analysis. Axial data represent the data with direction but no polarity, and angles $\theta$ and $\theta + \pi$ are treated as the same. Images of the mole crickets were measured as reflected images on a mirror placed under the experimental field. Therefore, the body angle of the mole crickets obtained here was equal to the burrowing trajectory, as seen from above in the experimental field.

(b) Mean axial vector of 24 individuals (Figure S3 (c)): In this experiment, three trials per individual were conducted for the burrowing experiments. For each of the 24 individuals, the mean body angle was calculated first, and the mean axial vector of the 24 individuals was calculated from this angle. To obtain the average body angle of each individual, the average angle was first calculated by doubling the body angles of all three trials per individual and the resulting angle was divided by two. This was plotted as the mean body angle of one individual on a circle of radius 1 (black points). From the 24 points plotted on the circumference, the mean axis vectors of the 24 individuals (red arrow) were calculated in a similar manner. Double arrows indicate the vectors. To indicate that the body angles are axial data, the points on the circumference are also indicated at 180° inverted positions.

Circular statistics (Rayleigh test for significant deviation from a random distribution and Watson's two-sample test for the homogeneity of two angular distributions) were conducted using the 'circular' package in R [1].

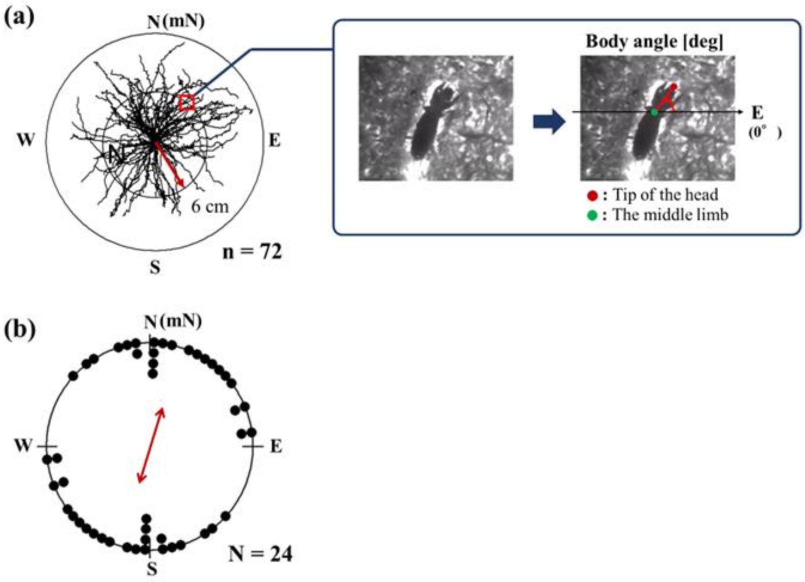


**Figure S3** Analysis of body orientation. (a) Extraction of body angles (e.g., 6 cm). From the burrowing trajectory of the mole crickets, the body angles of 72 trials at 6 cm from the starting point were calculated. The angle of the line connecting the tip of the head (red) and the middle of the mid-leg (green) was obtained from the images of mole crickets at that point (0° = north in the magnetic field in the laboratory (N)). (c) The mean axial vector (red arrow) for 24 individuals.

#### 3. Burrowing experiment with an inclination angle of the geomagnetic field

Unlike the outdoor geomagnetic field, the magnetic field inside the laboratory (LMF) has a small inclination angle. Therefore, we generated a magnetic field with the geomagnetic parameters of the ground (see Table S2, geomagnetic field (ground)) and conducted a burrowing experiment to examine whether inclination affected the burrowing behaviour of mole crickets. In this experiment, we used 19 mol crickets and performed three trials per individual for a total of 57 trials.

The burrowing trajectory (Figure S4(a)) for the 57 trials under the ground geomagnetic conditions resulted in a bimodal trajectory extending in the north-south direction from the centre. The distribution of the 19 individuals plotted in the mean body angle data of one individual was also significantly oriented at all three measurement positions (Rayleigh test: $P < 0.05$), and the mean vector (red arrow) was oriented in a north-south direction (Figure S4(b)).

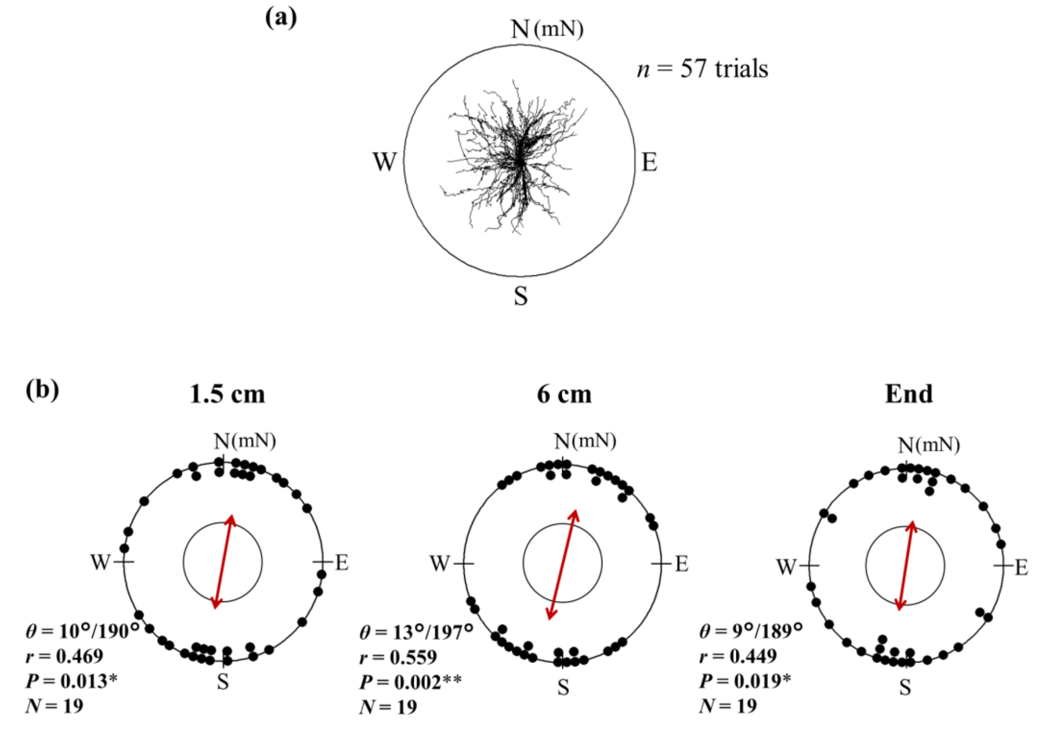


**Figure S4 Burrowing behaviour with an inclination angle comparable to the geomagnetic field.** (a) Burrowing trajectories for all 57 trials. (b) The mean axial vector of 19 individuals at three measurement points. The plotted points represent the mean body angle of one individual, from which the mean axis vector (red arrow) of the 19 individuals was calculated. N, E, S, and W around the circumference represent the azimuth in the magnetic field in the laboratory , and mN represents the magnetic north, respectively. *n* and *N* represent the numbers of trials and individuals, respectively. The inner circles in (b) and (c) represent the significance level of $P = 0.05$ calculated using the Rayleigh test; $\theta$ is the angle of the mean axis vector when N is 0°, r is the length of the mean axis vector, and *P* is the *P*-value calculated with the Rayleigh test.

**Supplemental reference**

[1] Agostinelli, C. & Lund, U. 2022 R package: Circular Statistics (version 0.4-95).
